# A slow-exchange conformational switch regulates off-target cleavage by high-fidelity Cas9

**DOI:** 10.1101/2020.12.06.413757

**Authors:** Viviane S. De Paula, Abhinav Dubey, Haribabu Arthanari, Nikolaos G. Sgourakis

**Affiliations:** Department of Pathology and Laboratory Medicine, The Children’s Hospital of Philadelphia, and Department of Biochemistry and Biophysics, Perelman School of Medicine, University of Pennsylvania, 3401 Civic Center Blvd, Philadelphia, PA, 19104, USA; Núcleo Multidisciplinar de Pesquisa em Biologia, Universidade Federal do Rio de Janeiro, Duque de Caxias, RJ, 25245-390, Brazil; Department of Biological Chemistry and Molecular Pharmacology, Harvard Medical School, and Department of Cancer Biology, Dana-Farber Cancer Institute, Boston, MA 02215, USA

## Abstract

The Cas9 endonuclease is broadly used for genome engineering applications by programming its single-guide RNA, where high specificity is required. Although mechanistic understanding of DNA cleavage in the CRISPR-Cas9 system has been guided by crystallographic structures and single-molecule FRET experiments, the role of conformational plasticity within its DNA recognition lobe (REC) for target accuracy remains elusive. Through NMR analysis of milliseconds-timescale dynamics, we show that the REC3 domain exchanges between a major, closed and a minor, open conformation. We find that a single mutation in the HiFi Cas9 variant (R691A) increases conformational dynamics in REC3, eliciting a global transition to the open conformation which is regulated by an intramolecular salt-bridge network. These observations suggest a mechanism for reduced off-target recognition by switching on a global conformational exchange process to allow REC3 scanning for proper base pairing. Our data establish a framework for rational engineering Cas9 variants with improved target discrimination.

## Introduction

The clustered regularly interspaced short palindromic repeats (CRISPR)-associated endonuclease Cas9 from *Streptococcus pyogenes* (SpyCas9) has revolutionized precise genome editing applications in basic research and represents an enormous promise for the development of novel therapeutic approaches for human diseases (*1*–*4*). However, a main concern for the widespread adoption of the CRISPR-Cas9 technology are unintended off-target events which occur when Cas9 cleaves a mismatched DNA/RNA heteroduplex at the distal end to the protospacer adjacent motif (PAM) (*5*–*7*).

To enhance the specificity of target cleavage, several rationally designed and engineered *S. pyogenes* Cas9 variants have been developed, leading to significant specificity improvements in unbiased genome-wide CRISPR-Cas9 measurements, comprising: high-fidelity (SpCas9-HF1), enhanced specificity (eSpCas9), hyper-accurate (HypaCas9), evolved (evoCas9) and high-fidelity (HiFi Cas9) (*8*–*13*). Notably, all of these variants involve amino acid substitutions of the non-catalytic REC3 domain within the larger DNA recognition lobe (REC), which has been shown to provide a template proofreading function by recognizing target complementarity and triggering long-range activation of the HNH nuclease domain (*11*). However, the mechanism by which correct base pairing at REC3 triggers the transition of the HNH domain from the inactive to the active state remains unknown. A crystal structure with mismatches at the PAM-distal end bound to a high-fidelity enzyme is not available, consequently important data to define the molecular basis for specificity of high-fidelity variants are lacking. Single-molecule fluorescence resonance energy transfer (smFRET) measurements and a detailed single turnover kinetic analysis showed that high-fidelity Cas9 variants increase discrimination largely by slowing the observed rate of cleavage without increasing the rate of DNA rewinding, favoring the release rather than cleavage of a bound off-target substrate (*14*, *15*).

In the current work, we use solution-state nuclear magnetic resonance (NMR) spectroscopy focusing on highly sensitive methyl groups as functional probes to show that the REC3 domain exists in a dynamic equilibrium between a ground and a sparsely populated state which interconvert on the milliseconds timescale. We used our system to characterize a high-fidelity REC3 mutant (R691A) (*13*) and found a drastic change in conformational exchange, which shifts the equilibrium toward the excited-state conformation. Careful inspection of X-ray structures suggested that the conformational transition between the two states is regulated by an intramolecular network of salt-bridges impacting the conformation of the α30-α32 helical motif. Based on our findings, we propose that HiFi Cas9 achieves its improved specificity by slowing down the conformational exchange process in the REC3 domain, which ultimately leads to a more stringent conformational proofreading for mismatches in the DNA target. Insights derived from our data underline the molecular basis for HiFi Cas9-mediated sensing of PAM-distal end mismatches, and establish a framework to further improve the specificity of Cas9 for genome engineering applications.

## Results

### A local conformational exchange process in the REC3 domain revealed by methyl NMR

SpCas9 is a 160 kDa multi-domain nuclease whose size challenges traditional solution-state NMR approaches for studies of protein structure and dynamics. Herein, as in our previous study of Cas9 (*Nerli and De Paula, 2020, under revision*), we have used a “divide-and-conquer” strategy to design optimized constructs of individual domains for investigating the dynamics of Cas9 by NMR (**Fig. 1**). We have exploited stereospecific methyl labeling of highly deuterated protein domains in concert with our recent automated methyl resonance assignment method to obtain well-resolved, high-quality spectra and complete methyl assignments for several Cas9 domains (*Nerli and De Paula, 2020, under revision*). To gain insights into conformational changes on Cas9 domains in solution, we focused on a 25 kDa construct of REC3 domain (WT and mutant), a subdomain of the REC lobe that was prepared as a highly deuterated, selectively Ile ^13^Cδ_1_; Leu ^13^Cδ_1_/^13^Cδ_2_; Val ^13^Cγ_1_/^13^Cγ_2_-labeled protein sample (**Fig. 1c**). The resulting protein showed well-dispersed 2D NMR spectra, indicating stable, properly conformed monomeric REC3, free of aggregation or degradation, as characterized by size exclusion chromatography coupled to multi-angle laser light scattering (SEC-MALS) (**Figure 1– figure supplement 1**).

**Figure 1.**
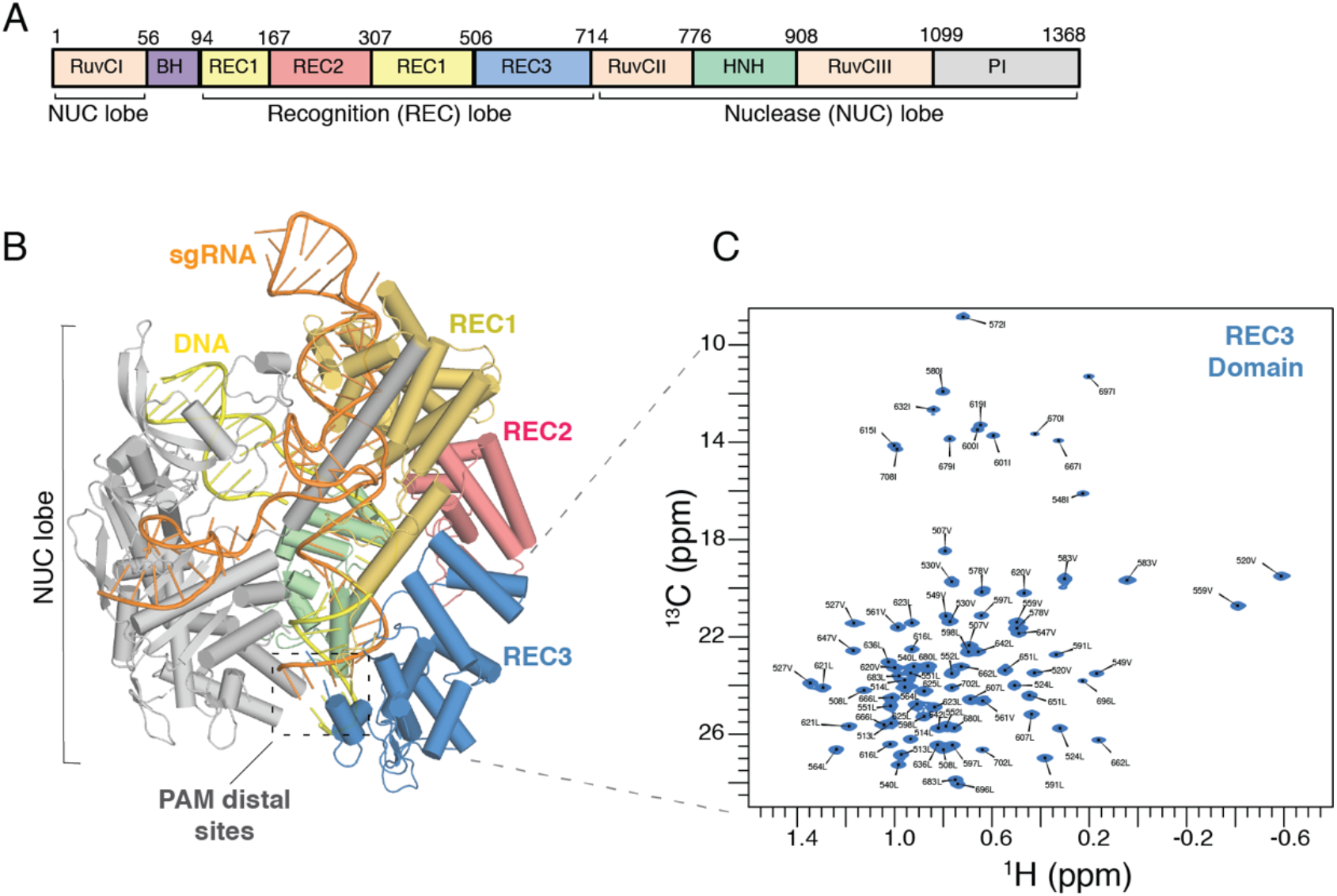
Domain structure and methyl resonance assignments of the REC3 Cas9 domain. Domain organization of *S. pyogenes* Cas9 composed of the recognition lobe (REC) and nuclease lobe (NUC). BH, bridge helix; PI, PAM-interacting. (B) Crystal structure of sgRNA/DNA-bound Cas9 (Protein Data Bank (PDB) accession number 4UN3) (*18*). Cas9 is shown in cartoon, with protein domains in different colors. The X-ray structure captures the inactive state of the HNH domain, which is a conformational checkpoint between DNA binding, and cleavage. The dotted black box highlights the location of base pair mismatches associated with off-target effects, at PAM distal sites. (C) ^1^H,^13^C-HMQC spectra of selectively labeled REC3 domain at Ile δ_1_-^13^CH_3_; Leu, Val-^13^CH_3_/^13^CH_3_ positions, acquired at 800 MHz, 25 °C (Nerli and De Paula, 2020).

To characterize the conformational landscape sampled by REC3 in solution, we performed a series of ^13^C single-quantum methyl Carr-Purcell-Meiboom-Gill (CPMG) relaxation dispersion experiments (*16*). Our data can detect conformational exchange processes that are active on the milliseconds-to-microseconds timescale, impacting the local magnetic environment of specific methyl groups distributed throughout the REC3 domain structure. Briefly, in CPMG experiments the effective transverse relaxation rate (*R*_2,eff_) is measured as a function of refocusing pulse frequency (νCPMG) which quenches the effects of conformational exchange, producing so-called dispersion profiles. Dispersion profiles, *R*_2,eff_ vs. νCPMG, where *R*_2,eff_ decreases with νCPMG, provide a hallmark of chemical exchange and, when fit to an appropriate model of exchange, estimates of the excited-state population (*p*_E_), interconversion rate (*k*_ex_) and chemical shift differences between the two exchanging states |Δω| (ppm) are obtained. In contrast, flat dispersion profiles suggest an absence of milliseconds-timescale dynamics. Here, we observe marked exchange contributions (> 5 s^−1^), as illustrated for representative methyl groups (L666δ_2_, L683δ_1_, I679δ_1_), showing that REC3 is undergoing a local conformational exchange processes on the milliseconds-to-microseconds timescale (**Fig. 2a**). The remaining methyl resonances exhibit flat dispersion curves or are in regions of high spectral overlap, prohibiting quantitative analysis. The dispersion curves for 6 methyl resonances can be captured by a simple two-site exchange model, to obtain a *k*_ex_ value of 988 ± 88 s^−1^ and a *p*_E_ value of 0.8 ± 0.05% (**Fig. 2b and Figure 2– figure supplement 1**). The methyl groups with marked exchange contributions are all located at the hydrophobic interface between the α30, α31 and α32 α-helices of the REC3 domain (PDB ID 4UN3) (**Fig. 2c**) and form direct contacts with nucleic acid residues at the PAM-distal end in the full Cas9 structure (**Figure 2– figure supplement 2.**). This region has been shown to undergo significant structural changes upon RNA/DNA heteroduplex recognition (*11*, *17*–*19*), suggesting that the observed exchange processes sampled by free REC3 in solution may help promote the crystallographically observed conformational changes associated with target recognition to regulate overall catalytic competence. Taken together, our current solution NMR data in conjunction with previous structural studies suggest that conformational plasticity of the α30-α32 REC3 helical motif is a key component of the solution dynamics of WT Cas9, relevant for the formation of a specific RNA-DNA heteroduplex.

**Figure 2.**
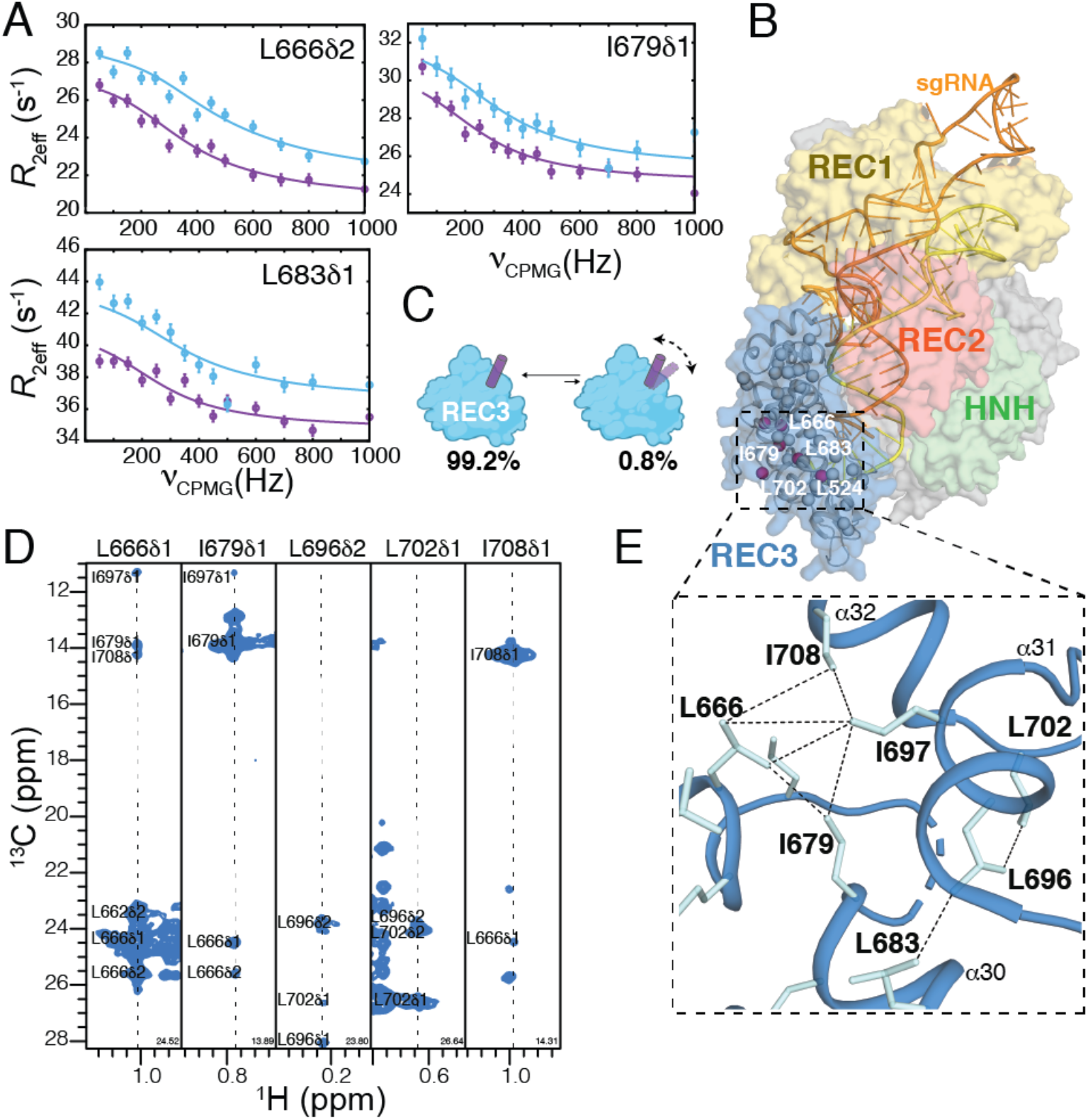
Conformational dynamics and ground-state structure of Cas9 REC3 domain. (A) Methyl-selective ^13^C single quantum CPMG relaxation dispersion profiles carried out at two magnetic fields (600 MHz, purple; 800 MHz, blue) are shown for selected residues (L666δ_2_, I679δ_1_ and L683δ_1_) of WT REC3 domain (25 kDa). Experimental data are shown as small circles in all panels, with errors estimated from the S/N in the raw experimental data. The best-fit lines are shown for a global analysis of the resonances of 6 methyl groups exhibiting significant relaxation dispersion profiles using a two-site conformational exchange model. (B) Architecture of the Cas9 protein (PDB ID 4UN3) highlighting its protein domains as follow: REC1 (yellow), REC2 (red), REC3 (blue), HNH (green) and RuvC (gray). Six methyl probes undergoing chemical exchange by CPMG are shown as spheres on the REC3 domain and colored purple. (C) Schematic representation of REC3 domain illustrating the interconversion between a ground state, which is ~99.2% populated, and an invisible, excited state, which is ~0.8% populated. The doted arrows represent the conformational transition of the alpha-helix 31 (α31). (D) ^13^C_M_-^1^H_M_ strips from a 3D C_M_-C_M_H_M_ SOFAST NOESY experiment taken at the ^13^C_M_ coordinates of stereospecifically assigned methyl resonances noted on each panel, showing NOE cross-peaks between the methyl resonances of residues on helices α30, α31 and α32. (E) Close-up view of the α30, α31 and α32 regions showing the network of observed NOEs (black dotted lines), corresponding to the major (ground-state) solution conformation.

To characterize the major (ground-state) conformation sampled by the α30, α31 and α32 helices of REC3 in solution, we analyzed methyl NOE intensities recorded in a 3D C_M_-C_M_H_M_ SOFAST NOESY experiment (**Fig. 2d**), relative to the corresponding features observed in *i*) the Cas9 apo structure (PDB ID 4CMP) and *ii*) the Cas9-sgRNA-DNA ternary complex (PDB ID 5Y36). Consistently with the Cas9 structure in the apo state, we find that the network of observed short-range NOEs connecting the methyl groups of L666, I679, L683, L696, I697, L702, L708 located at the vicinity of the α30, α31 and α32 helices, (**Fig. 2e, black dotted lines**) is consistent with a “closed”, well-packed conformation where the sidechains of L666, I697 and L708 are forming direct hydrophobic contacts. This is evident from the observation of several unambiguously assigned short-range (3.5 - 5 Å upper limit) NOEs between the corresponding methyl groups (**Fig. 2d, e**). On the contrary, in the sgRNA-DNA ternary complex the helices α30, α31 and α32 are found in an “open” conformation, with the methyl groups of L666, I708 oriented at distances from the core methyls (~ 12 Å), well beyond the NOE detection limit of 10 Å (**source data 1**).

### Impact of mismatches on binding and cleavage by HiFi Cas9

We wanted to determine how mismatches between guide RNA and target DNA influence the binding and cleavage by Cas9 mutants. We focused on the high fidelity Cas9 (HiFi Cas9) (*13*), a Cas9 variant containing a single point mutation (R691A) identified by an unbiased bacterial screening approach, which showed reduced global off-target effects while maintain high on-target activity when used as a ribonucleoprotein (RNP) complex. When compared to the rationally designed eSpCas9(1.1), SpCas9-HF1, and HypaCas9 high-fidelity Cas9 mutants, HiFi Cas9 (R691A), also exhibited reduced on target editing at many sites when used as a RNP (*13*). We used a fluorescence anisotropy (FA) assay to test the binding of DNA sequences that contain pairs of substituted bases at positions ranging from 17 to 20 (numbering starting with 1 for the most PAM-proximal base and ending with 20 for the most PAM-distal base) (**Fig. 3a, b and source data 2**). Consistent with previous binding measurements for several Cas9 variants (eSpCas9(1.1), HypaCas9 and Cas9-HF1) (*11*), the affinities (*K*_d_ ~20 nM) of HiFi Cas9 (R691A) for on-target and PAM-distal mismatched DNA substrates were similar to WT Cas9 (**Fig. 3c**), demonstrating that cleavage specificity is enhanced through a mechanism distinct from a reduction of target binding affinity.

**Figure 3.**
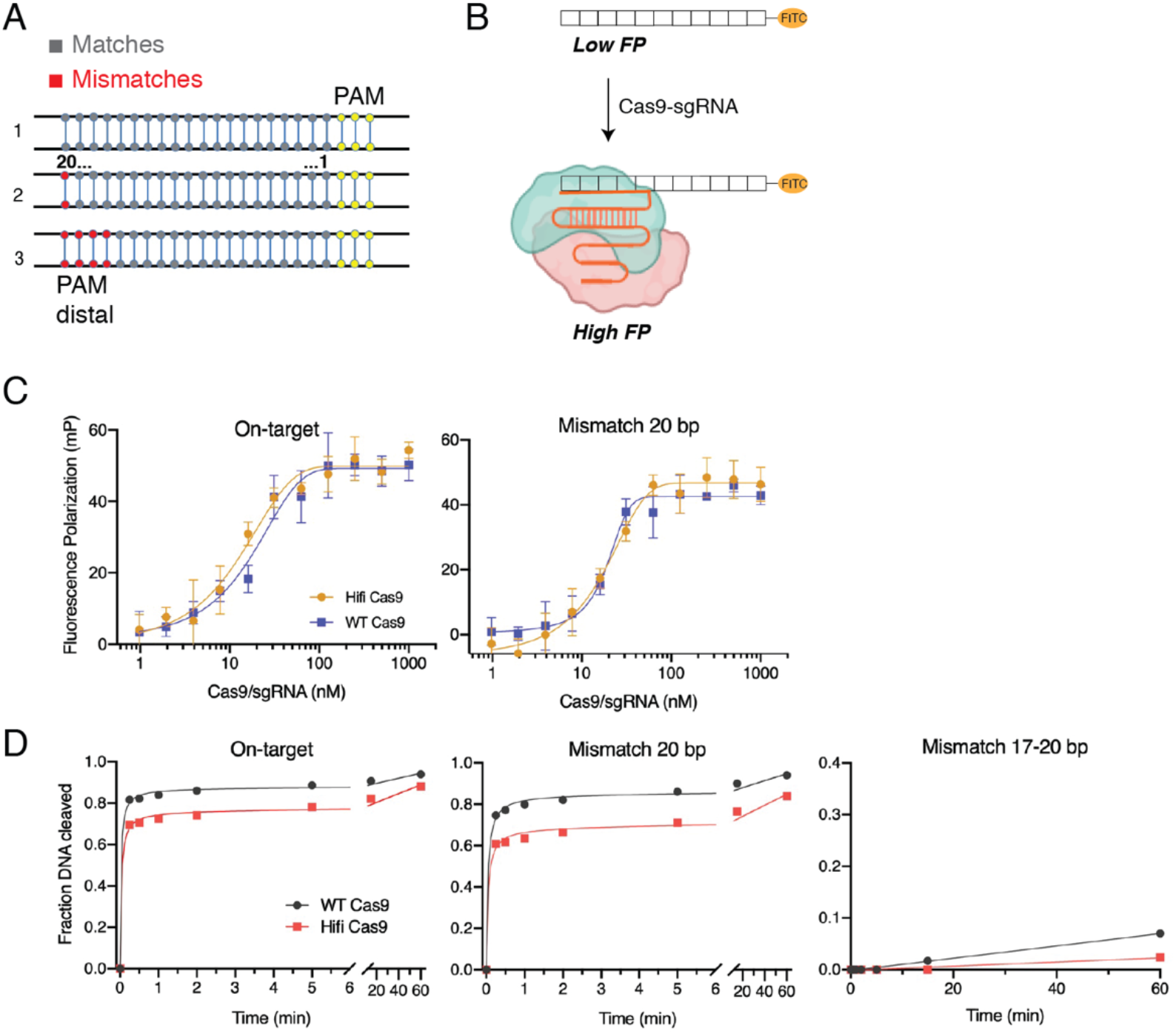
Influence of mismatches between the sgRNA and target DNA on binding and cleavage by HiFi-Cas9 R691A. (A) Scheme of the dsDNA substrates with on-target and mismatches in the protospacer distal region (PAM distal) against the guide RNA used in the *in vitro* assays. The number of mismatches in the PAM-distal regions are shown in red. (B) Schematic representation of a fluorescence polarization (FP)-based assay for monitoring Cas9:gRNA and DNA binding. (C) Dose-dependent increase in the FP signal (in millipolarization, mP units) upon binding of on-target and mismatched DNA to the Cas9:gRNA complex. Error bars represent ±1 SD across technical replicates (n = 3). (D) Quantification of DNA cleavage assay time course of WT Cas9 versus HiFi Cas9 on perfect and PAM-distal mismatched targets. Exponential fits are shown as solid lines.

To evaluate how mismatches between sgRNA and target DNA influence the cleavage by HiFi Cas9, we performed an *in vitro* kinetic cleavage assay which monitors the cleavage of the *EMX1* target sequence with or without nucleotide mismatches to the guide RNA. Consistently with previous assays reported for other Cas9 variants (*11*), alanine substitution of Arg691 yielded an enzyme with moderately reduced cleavage activities towards linearized plasmid DNA containing a perfectly complementary sequence, but almost no activity towards DNA containing 4 mismatches to the guide RNA at positions 17-20 (**Fig. 3d and Figure 3– figure supplement 1**). A reduction in global off-target editing for HiFi Cas9 while maintaining high on-target activity when used as a ribonucleoprotein delivery complex has been described previously (*13*). Together, these results establish the enhanced mismatch intolerance of HiFi Cas9 using our recombinant *in vitro* system.

### The REC3 HiFi Cas9 mutant exhibits altered dynamics of a helical interface region

We next sought to investigate the structural mechanism underlying the improved discrimination against off-target DNA with mismatches at the distal end of the guide RNA in the HiFi Cas9 (R691A) variant. To study the solution structure and conformational dynamics of the HiFi Cas9 variant, we prepared a selective ^1^H/^13^C ILV (Ile, Leu and Val)-methyl labelled R691A REC3 sample, and compared the ^13^C-^1^H SOFAST HMQCs spectra between the wild type REC3 and the R691A mutant. Due to the exquisite sensitivity of methyl chemical shifts to structural perturbations, a rapid assessment of the effects of mutations on protein structure can be obtained through comparison of ^1^H-^13^C HMQC spectra of WT and mutant REC3. Significant chemical shift differences (greater than 1 s.d. [0.05 parts per million (ppm)] or in the range 0.5 to 0.1 ppm relative to the ^13^C field) were observed for the methyl resonances of several residues, indicating that the R691A REC3 mutation, located on helix α31, induces effects impacting the entire α30-α32 helical motif as well as methyl groups located on the distal part of the domain, as far as 26 Å away from the mutation site (**Figure 4– figure supplement 1**). In addition, while in the WT REC3 domain most methyl groups give rise to well-resolved NMR peaks, in the R691A mutant several resonances are broadened (corresponding to the methyl groups of 13 residues), likely due to increased conformational exchange at the intermediate (microsecond to millisecond) NMR timescale (**Figure 1– figure supplement 1 and Figure 4– figure supplement 1**). These observations suggest that the R691A mutation induces a global perturbation in dynamics of the REC3 domain.

**Figure 4.**
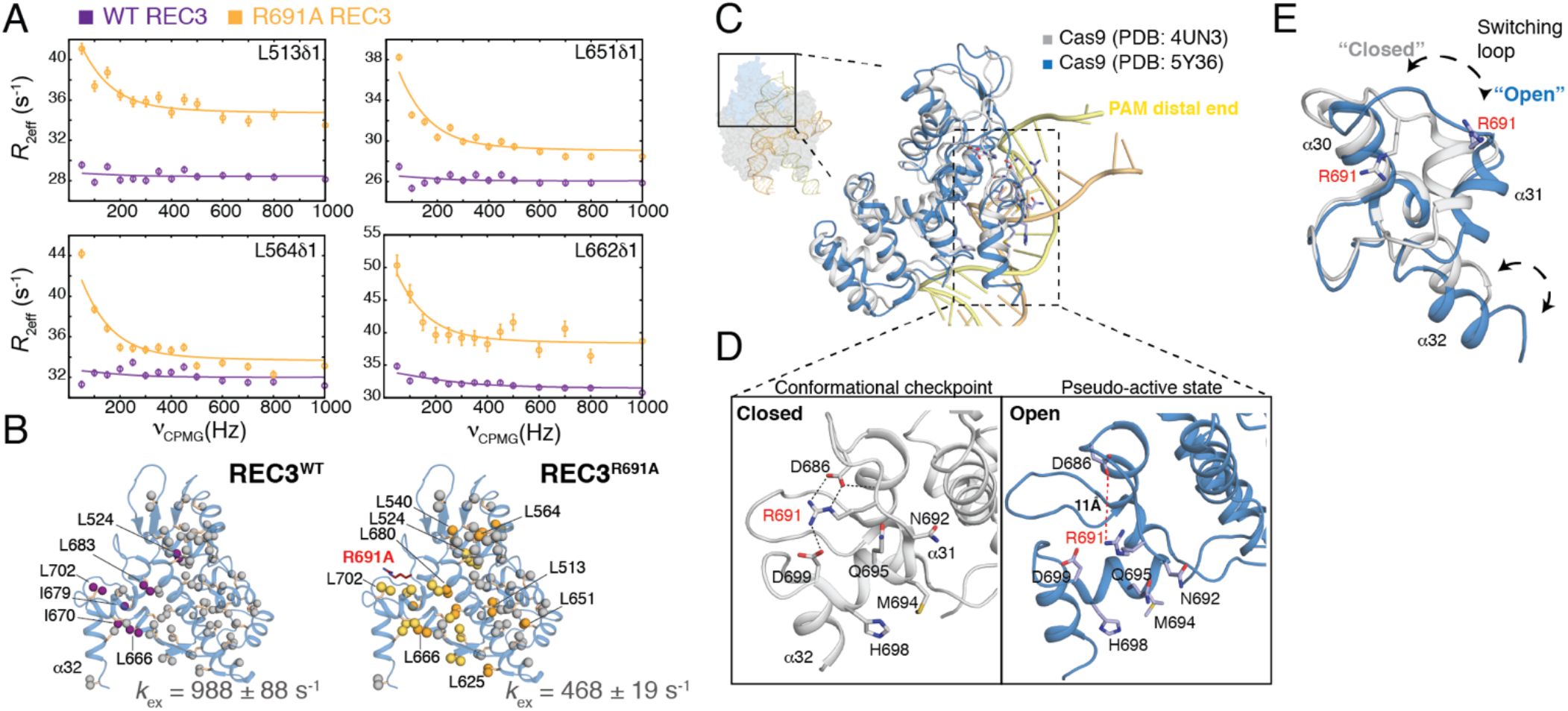
A REC3 salt-bridge switch defines the conformational plasticity of high-fidelity Cas9. (A) Comparison of methyl ^13^C-CPMG relaxation dispersion profiles for L513δ_1_, L564δ_1_, L651δ_1_ and L662δ_1_ of WT REC3 (purple) and R691A REC3 (orange), at 600 MHz and 15 °C. (B) Residues that show an exchange contribution, *R*_ex_ > 5 s^−1^ are shown as purple (WT REC3) and orange (R691A REC3) spheres. Residues with line broadening beyond detection are colored in yellow and residues with no dispersion are shown in gray. Fitted *k*_ex_ values from dispersion experiments are shown next to the structure, with χ^2^ fit surfaces shown in **Figure 2– figure supplement 1**. (C) Superposition of the overall structures of Cas9/sgRNA/DNA (PDB: 4UN3, in gray and PDB: 5Y36, in blue) and zoomed-in view of the REC3 domain. (D) Close-up view of REC3 domain sidechain contacts in the conformational checkpoint and pseudo-active Cas9-nucleic acid complex structures. The α31 helix in REC3 adopts either a closed (checkpoint state) or an open conformation (pseudo-active state). The closed conformation is stabilized by salt-bridge interactions between the side chain of R691 and the side chains of D686 and D699 (black dashes). (E) Distinct opening-closing motions of the α31 helix highlighting the transition of the REC3 domain from the conformational checkpoint to the pseudo-active state.

Since it is known that REC3 domain forms direct contacts with the RNA-DNA heteroduplex and undergoes long-range conformational changes that are important for target recognition (*11*, *20*, *21*), we aimed to characterize changes in dynamics introduced by the R691A HiFi mutant. We therefore performed methyl ^13^C-CPMG relaxation dispersion experiments under identical sample concentration and other conditions, and compared the resulting relaxation dispersion data to our established results for the WT form. We observe that the exchange contributions to the ^13^C-CPMG relaxation rates show a marked increase in the R691A mutant, for methyl groups distributed throughout the REC3 structure (**Fig. 4 and Figure 4– figure supplement 2**). Data fitting of the CPMG relaxation curves show that the exchange process becomes 2-fold slower relative to the WT (*k*_ex_ of 468 ± 19 s^−1^) and the excited-state population increases significantly (68 ±14%, compared with 0.8 ±0.05% in the WT REC3), consistently with the observed chemical shift differences (**Fig. 4a, b** and **source data 3).** It is worth noting that our CPMG data can be fit using an excited-state population in the 15-70 % range with a modest decrease in fit quality relative to the global minimum, as shown by a detailed χ^2^ surface analysis **(Figure 2– figure supplement 1**). These observations suggest the presence of a global conformational exchange process on the μs-ms timescale, involving a movement of the α31 helix, and that the R691A mutation shifts the conformational equilibrium to stabilize the *excited* state.

Close inspection of X-ray structures of Cas9-sgRNA-DNA ternary complexes (*20*, *23*–*25*) reveal that, in the conformational checkpoint (*18*) (denoted as *closed state*), the α31 helix including the R691 establishes a network of salt-bridges with D686 and D699 and maintains its stability (**Fig. 4c)**. Conversely, a distinct conformational state was captured in a cryo-EM structure of dCas9-sgRNA-DNA, revealing that the R691 side-chain is oriented towards the solvent away from the D686 and D699 side chains (11 Å distance), and does not form any sidechain contacts with other residues (**Fig. 4d**). Additionally, EM-based MD simulations support that the cryo-EM structure represents the pseudo-active state (denoted as *open-state*) of SpCas9 in an environment primed for DNA cleavage (*22*). In the context of these structures, our NMR results support that in solution the R691 side chain helps to stabilize a closed, ground-state conformation of the REC3 domain by making several contacts within the α30-α31 helical region. Disruption of the conserved salt bridge network (**Figure 4–figure supplement 3)** in the R691A HiFi mutant shifts the conformational equilibrium toward the open, excited-state to induce a more global conformational exchange process occurring on a 2-fold longer timescale, relative to the WT (**Fig. 4e**). Overall, our current NMR data provide evidence that a loss of loop constraints in the REC3 domain led to a gain of dynamics.

## Discussion

On the basis of our solution NMR results, the REC3 domain exists in a conformational equilibrium between two conformations, corresponding to closed (inactive) and open (active) states. In the HiFi Cas9 mutant, this equilibrium is shifted toward the open state (*p*_*E*_ = 68 ±14%), by slowing down the conformational exchange rate between these two states. Considering that the closed state is the predominant state of the wild-type REC3 under physiological conditions, it is notable that the introduction of a single-point mutation, R691A, significantly alters the energy landscape to re-weight the relative populations of the two conformations. The NMR CPMG dispersion data support that the milliseconds-timescale dynamics originate from exchange between inactive and active conformers because significant differences in *p*_*E*_ were obtained. Furthermore, our methyl-based NMR measurements provide evidence that the R691 sidechain contacts with D868 and D699 are important for the conformational sensing of PAM-distal mismatches observed in the HiFi Cas9 variant. However, we cannot definitively rule out that the mutant introduces a completely novel conformational state, not sampled by WT REC3, without further investigation.

Analysis of our NMR and previous functional data in the context of the available Cas9 co-crystal structures suggests a general scheme for RNA-DNA heteroduplex target site recognition and cleavage (**Fig. 5**). We propose that the ground- to excited-state transition described here provides a mechanism to achieve an active, catalytically competent Cas9 state, promoting high accuracy DNA targeting. Stabilization of the excited state in HiFi Cas9 may work as a *fidelity check point* for DNA proofreading, leading to an enhanced scanning and stringent specificity for mismatches at the PAM-distal end pairing. Based on our analysis of crystallographically observed snapshots of Cas9-sgRNA-DNA complexes, we hypothesize that the excited-state conformation might share similar features with the catalytically competent state reported for the cryo-EM of dCas9-sgRNA-DNA structure (*22*).

**Figure 5.**
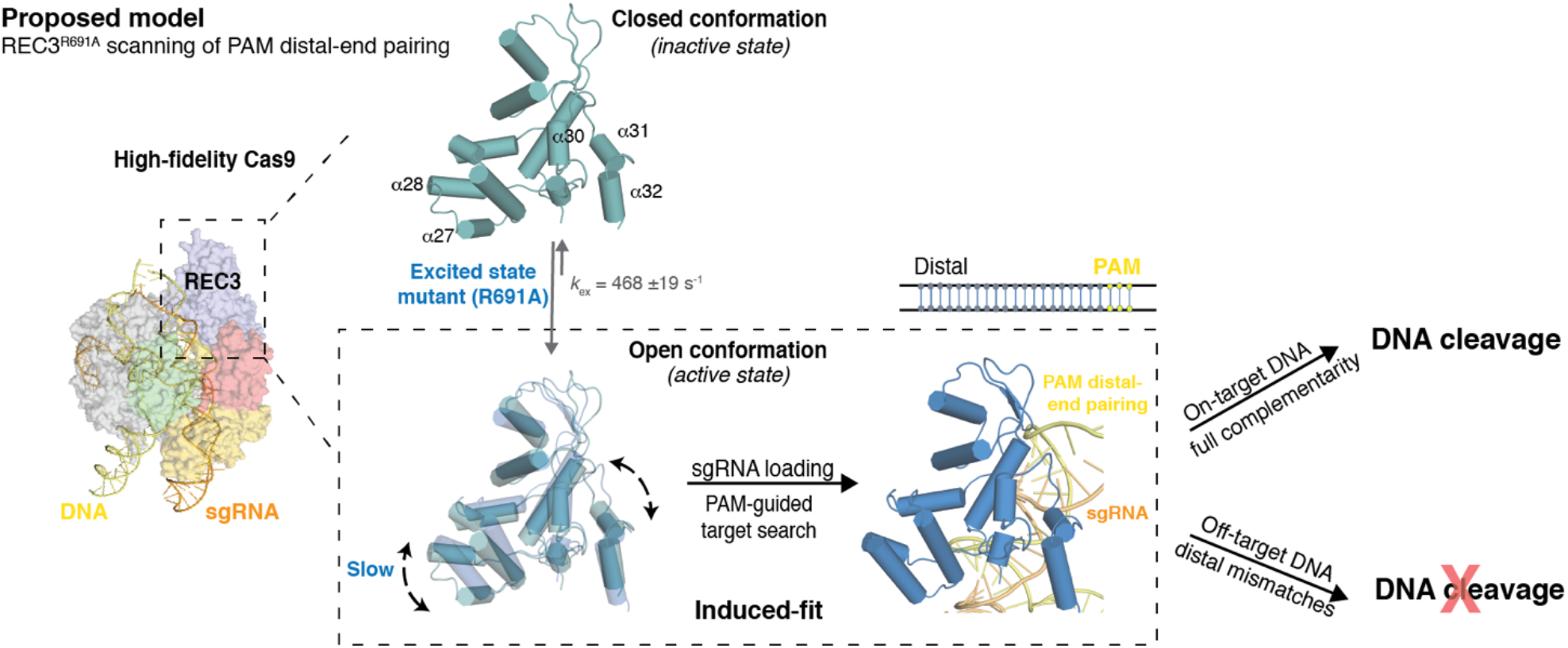
Conformational sensing by HiFi REC3-R691A drives PAM distal-end pairing specificity. Proposed mechanism for HiFi REC3 domain sensing and regulation of the RNA-DNA heteroduplex for allosteric HNH activation and cleavage. A global conformational transition enables REC3 to sample two distinct states. In the closed conformation, the R691 on the α31 helix forms a network of salt-bridges with D686 and D699. Disruption of the network by the R691A mutation shifts the equilibrium toward the open conformation (active state), inducing global changes in core dynamics and slowing mobility of the α31 helix, which ultimately lead to a stringent conformational proofreading for mismatches at the PAM distal-end in the DNA target.

Our data show that conformational dynamics make important contributions to the high fidelity Cas9, and the slow rate of the conformational transition from close to open state is the key determinant of enzyme fidelity in the present system. Evidence for a functional role of such slow conformational rates was provided in previous studies focusing on both engineered HypaCas9 and Cas9-HF1 enzymes, which demonstrated that high-fidelity Cas9 variants achieved improved discrimination against PAM-distal mismatches by reducing the observed rate of cleavage for off-target DNA as compared to WT Cas9, which shifts kinetic partitioning to favor release rather than cleavage of the bound DNA (*14*, *15*, *26*, *27*). While the open conformation mediates high-affinity complex formation with target DNA, the binding process and cleavage is likely achieved via the formation of an initial encounter complex, where additional induced structural adaptations follow the selection of pre-existing conformational states. An analogous induced-fit mechanism is used by DNA polymerase (*27*), and is consistent with the majority of protein-ligand binding examples, where both processes contribute to complex formation (*28*) (**Fig. 5**). Our results are consistent with the hypothesis that WT REC3 undergoes a more local conformational process that is consistent with a repacking of core hydrophobic sidechains. On the contrary, the R691A mutant induces a more global, concerted conformational transition of the α30-α32 helices and core of the structure, allowing REC3 to sense whether or not nucleotides at the PAM-distal end are base-paired with the guide RNA. In contrast, a mismatched base fails to transduce the formation of the high-affinity DNA binding state, thus facilitating the structural transition to the closed state and release of the mismatched region. Thus, protein dynamics can facilitate an allosteric pathway that communicates REC3 to the catalytic HNH domain, leading to increased accuracy.

In summary, our study focusing on the REC3 domain highlights the potential of targeting Cas9 excited states to steer the conformational equilibrium towards states with maximum specificity. With ever-increasing interest in designing hyper-accurate Cas9 variants for therapeutic, diagnostic and biotechnology applications, our solution-based description of REC3 conformational dynamics, as well as similar studies focusing on other domains (*29*), can be leveraged to design CRISPR enzymes with properties that are tailored to specific applications.

## Materials and Methods

### *S. pyogenes* Cas9 REC3 domain selectively isotope labelling, NMR sample preparation and stereospecific methyl resonance assignments

Recombinant wild-type *Streptococcus pyogenes* Cas9 REC3 domain (506-712) possessing an amino-terminal His_10_-MBP tag (Addgene, no. 101205) followed by a TEV protease site was expressed in *Escherichia coli* strain BL21(DE3) and purified as described previously (*Nerli and De Paula*). U-[^15^N,^12^C,^2^H]-labelled REC3 domain was overexpressed in BL21(DE3) cells containing chaperone plasmid pG-KJE8 (TAKARA, 3340) in M9 medium in ^2^H_2_O containing 2 g l^−1^ ^2^H^12^C glucose (Sigma #552003) and 1 g l^−1^ ^15^NH_4_Cl. Selective ILV-methyl (Ile ^13^Cδ1; Leu ^13^Cδ1/^13^Cδ2; Val ^13^Cδ1/^13^Cδ2;) was achieved by the addition of appropriate precursors (ISOTEC Stable Isotope Products (Sigma-Aldrich) as detailed previously (*30*, *31*).

Briefly, when cells reached an OD_600_ of ~0.6, isopropyl β-d-1-thiogalactopyranoside (IPTG) was added to a final concentration of 0.5 mM to induce protein expression. Cells were then grown for an additional 18 h at 22 °C. Collected cells were resuspended in lysis buffer (50 mM Tris pH 7.5, 500 mM NaCl, 5% (v/v) glycerol and 1 mM Tris(2-carboxy-ethyl) phosphine (TCEP) containing an EDTA-free protease inhibitor tablet (Roche). The cell suspension was sonicated on ice and clarified by centrifugation at 27,000g for 15 min. The soluble lysate fraction was bound in batch to nickel-nitrilotriacetic acid (Ni-NTA) agarose (Qiagen). The resin was washed extensively with 20 mM Tris, pH 7.5, 500 mM NaCl, 1 mM TCEP, 10 mM imidazole, and 5% (vol/vol) glycerol; and the bound protein was eluted in 20 mM Tris, pH 7.5, 500 mM NaCl, 1 mM TCEP, 300 mM imidazole, and 5% (vol/vol) glycerol. The His_10_-MBP affinity tag was removed with His_6_-tagged TEV protease during overnight dialysis against 20 mM Tris, pH 7.5, 500 mM NaCl, 1 mM TCEP, and 5% (vol/vol) glycerol. The protein was then flowed over Ni-NTA agarose to remove TEV protease and the cleaved affinity tag. REC3 domain (WT and R691A mutant) were further purified by size-exclusion chromatography on a Superdex 200 16/60 column (GE Healthcare) in 20 mM Tris, pH 7.5, 200 mM KCl, 1 mM TCEP, and 5% (vol/vol) glycerol.

2D ^1^H-^13^C SOFAST-HMQC were acquired on 400 μM ILV-methyl-labelled WT and R691A REC3. The change in chemical shift (in p.p.m.) between the WT and R691A REC3 domain methyls was determined using the equation Δδ^CH3^ = [1/2 (Δδ^2^H + Δδ^2^C/4)]^1/2^. 3D C_M_-C_M_H_M_ SOFAST NOESY experiment was recorded at 800 MHz, 25 °C on ILV-methyl-labelled WT REC3 sample using a recycle delay of 0.2 s and NOE mixing time of 300 ms. Assigned NOEs were cross-validated based on the Cas9-sgRNA bound state structure (PDB ID 4ZT0) as described in (Nerli, De Paula, 2020). Both ILV labeled WT or R691A REC3 samples contained 400 μM protein in 20 mM sodium phosphate, pH 7.4, 200 mM KCl, 1 mM TCEP, and 5% (vol/vol) glycerol d-8, 0.01% NaN_3_, in 90% H_2_O/10% D_2_O. All spectra were processed with NMRPipe (*32*) and analyzed with CcpNMR program (*33*).

### Stereospecific isotopic labeling

A specifically methyl-labeled acetolactate precursor (2-[^13^CH_3_], 4-[^2^H_3_] acetolactate) was obtained through deprotection and exchange of the protons of the methyl group in position 4 of ethyl 2-hydroxy-2-(^13^C)methyl-3-oxobutanoate (FB reagents) achieved in D_2_O at pH 13 (*34*). Typically, 300 mg of ethyl 2-hydroxy-2-(^13^C)methyl-3-oxobutanoate was added to 24 mL of a 0.1 M NaOD/D_2_O solution. After 30 min, the solution was adjusted to neutral pH with DCl and 2 mL of 1 M Tris pH 8.0 in D_2_O was added. For the production of highly deuterated [U-^2^H], I-[^13^CH_3_]δ1, L-[^13^CH_3_]proS, V-[^13^CH_3_]proS WT REC3 samples, 300 mg/L of 2-[^13^CH_3_], 4-[^2^H_3_] acetolactate, prepared as described above, was added 1 h prior to induction (OD_600_ ≈ 0.55). 40 min later (i.e. 20 min prior to induction), 3,3-[^2^H_2_],4-[^13^C]-2-ketobutyrate (SIGMA #589276) was added to a final concentration of 60 mg/L. Protein was induced at OD_600_ ≈ 0.7 by addition of 1 mM IPTG and expression was performed for 20 h at 23 °C.

### Expression and purification of Cas9

Recombinant wild-type and mutant *Sp*Cas9 (1-1,368) possessing a C-terminal decahistidine tag (Addgene, no. 62731) was expressed and purified as described previously (*35*). Proteins were expressed in *Escherichia coli* strain BL21(DE3) cells containing chaperone plasmid pG-KJE8 (TAKARA, 3340). When the optical density at 600 nm reached 0.6, protein expression was induced with 0.5 mM IPTG, and the temperature was lowered to 20 °C. The induction was then continued for 16-18 h. Collected cells were resuspended in lysis buffer (50 mM Tris pH 7.5, 500 mM NaCl, 5% (v/v) glycerol and 1 mM TCEP containing an EDTA-free protease inhibitor tablet (Roche). The cell suspension was sonicated on ice and clarified by centrifugation at 27,000g for 15 min. The soluble lysate fraction was bound in batch to nickel-nitrilotriacetic acid (Ni-NTA) agarose (Qiagen). The resin was washed extensively with 20 mM Tris, pH 7.5, 500 mM NaCl, 1 mM TCEP, 10 mM imidazole, and 5% (vol/vol) glycerol, and the bound protein was eluted in 20 mM Tris, pH 7.5, 500 mM NaCl, 1 mM TCEP, 300 mM imidazole, and 5% (vol/vol) glycerol. Fractions containing Cas9 were dialyzed into buffer A (20 mM Tris-Cl, pH 7.5, 125 mM KCl, 5% glycerol, 1 mM TCEP) and applied it onto a 1 ml HiTrap SP HP Sepharose column (GE Healthcare). Cas9 was eluted using a linear gradient from 0-100% buffer B (20 mM Tris-Cl, pH 7.5, 1 M KCl, 5% glycerol, 1 mM TCEP) over 20 column volumes. The protein was further purified by gel-filtration chromatography on a Superdex 200 16/60 column (GE Healthcare) in Cas9 storage buffer (20 mM Tris-Cl, pH 7.5, 200 mM KCl, 5% glycerol, 1 mM TCEP). Cas9 was flash frozen in liquid nitrogen and stored at −80 °C.

### Site-directed mutagenesis

The protein variants, SpyCas9^R691A^ full-length and REC3^R691A^ domain were generated using site-directed mutagenesis on the Cas9 and MBP-REC3 plasmids, respectively, using the Phusion High-Fidelity DNA Polymerase (New England Biolabs). Template DNA was removed by Dpn I treatment and transformed into *E. coli* DH5α strain. The introduced mutations and the absence of secondary mutations were verified by sequencing of plasmid DNA. Overexpression and protein purification were carried out as described above.

### Size‑exclusion chromatography in line with multi‑angle light scattering (SEC‑MALS)

Absolute molecular weight calculations were obtained by multi angle light scattering coupled with refractive interferometric detection (Wyatt Technology Corporation) and a Superdex 75 10/300 column (GE Healthcare) at 25 °C. REC3 protein samples (injection volume of 100 ΔL at 5 mg/mL) were run at a 0.5 mL/min flow rate in a running buffer of 20 mM sodium phosphate (pH 7.4), 150 mM NaCl and 1 mM DTT. Protein concentrations were monitored by a refractometer and light scattering directly after the gel filtration column. Absolute molecular weights were determined using ASTRA version 6.0 (Wyatt Technologies).

### Methyl CPMG relaxation dispersion experiments

Methyl single-quantum ^13^C CPMG relaxation dispersion experiments (*16*) were recorded on highly deuterated ILV-methyl labeled WT and R691A REC3 samples (both at 400 μM protein concentration) at field strengths of 14.0 T and 18.8 T, at 15 °C, using Bruker spectrometer, equipped with a cryogenically cooled probe. The CPMG data set was acquired as pseudo 3D experiments with a constant relaxation time period T_relax_ of 20 ms and with 15 CPMG pulse frequencies ν_CPMG_ = 1/(2τ) ranging from 50 to 1000 Hz, where τ is the delay between the consecutive 180° refocusing pulses in ^13^C CPMG pulse-train. Relaxation dispersion profiles R_2,eff_(ν_CPMG_) were calculated from peak intensities (I) recorded at different CPMG frequencies ν_CPMG_ using the following equation: R_2,eff_(ν_CPMG_) = −1/T_relax_ln(I/I_0_), where I is signal intensity in the spectra collected at T_relax_ = 20 ms, I_0_ is signal intensity in the reference spectrum recorded at T_relax_ = 0. An interscan delay of 1.5 s was used with 16 scans/FID, giving rise to net acquisition times of 23 h for a complete pseudo-3D data set. All data were processed using NMRpipe (*32*) and peak intensities were picked using CCPN (*33*). The error was determined from the noise level of the spectra. The variation in R_2,eff_ with ν_CPMG_ was fit to a two-state model of chemical exchange based on the Bloch-McConnell equations, to extract values of exchange parameters (*p*_B_, *k*_ex_=*k*_AB_+*k*_BA_), as well as ^13^C chemical shift differences for nuclei interconverting between pairs of states. The software CATIA (*36*) was used to fit the data. Initially, global fits included 6 profiles for WT REC3 at two magnetic fields (L666δ2, L683δ1, L696δ1, I679δ1, L702δ1, L702δ2). The fitting was performed by minimizing the function χ^2^ as previously described (*37*). The group fit of selected residues was performed if the χ^2^_Group_/χ^2^_Local_ was less than 2.0. For R691A REC3, analysis included 13 profiles at two magnetic fields (L513δ1, L514δ1, I548δ1, L551δ1, V559δ2, L564δ1, L598δ2, L621δ1, L623δ1, L625δ2, L651δ1, L651δ2, L662δ1). χ^2^ error surface plots were generated for WT and R691A REC3 to evaluate the robustness of the extracted exchange parameters (*pb*, *k*_ex_). Numerical fitting was performed using the CATIA program as before, with *pb* and *k*_ex_ sampled from a grid with values ranging from 0-80% and 0-2000 s^−1^, respectively and |Δδ| was free to change during the χ^2^ minimization procedure.

### *In vitro* transcription and purification of EMX1 sgRNA

To produce plasmid template at a large-scale, maxi prep were done according to manufacturer’s instructions (Qiagen 12662) and pooled. 1.5 mg of pNL003 plasmid (*38*) was linearized by incubation at 37 °C for 2-4 h in the presence of EcoRV (NEB R0195S). Transcription reactions (10 mL) were conducted in buffer containing 50 mM Tris, pH 8.1, 25 mM MgCl_2_, 0.01% Triton X-100, 2 mM spermidine, and 10mM DTT, along with 5 mM each of ATP, GTP, CTP, and UTP, 100 µg/mL T7 polymerase (made in house), and ∼1 µM DNA template. Reactions were incubated at 37 °C overnight and subsequently treated with 5 units of DNase (Promega) for 1 hour. Precipitants were removed by centrifugation and the transcribed RNA ethanol precipitated. RNA was then solubilized in denaturing loading dye (2 mM Tris/Cl pH 7.5, 20 mM EDTA, 8 M Urea, 0.025% (w/v) bromophenol blue, 0.025% (w/v) xylene cyanol), and loaded onto a 10% (wt/vol) polyacrylamide gel (29:1 Acrylamide:Bis-acrylamide), 7 M urea, 0.5× TBE. The appropriate band was excised, crushed and passively eluted from the gel overnight in 0.3 M sodium acetate in diethylpyrocarbonate (DEPC)-treated water at room temperature. The RNA was pelleted by centrifugation at 4000 xg for 60 min. The pellet was resuspended in 1 mL 70% (v/v) ethanol and transferred to a 1.5 mL Eppendorf tube and repelleted for 20 min at 21,000 xg. The pellet was washed and repelleted a second time in 70% (v/v) ethanol then dissolved in DEPC-treated water. The purity of the EMX1 sgRNA was confirmed by electrophoresis on a 12% TBE-UREA-PAGE gel (Thermo Scientific). Concentrations were determined by A_260nm_ using a NanoDrop instrument (Thermo Scientific).

### *In vitro* DNA cleavage Assays

SpCas9 WT and HiFi R691A mutant activity was assessed in an *in vitro* DNA cleavage assay with a linearized substrate plasmid DNA containing on target and mismatches at PAM distal-end positions. Prior to RNP formation, sgRNA EMX1 was refolded as previously described (*38*). Cas9 and refolded sgRNA were pre-incubated at room temperature for at least 10 min in 1× binding buffer (20 mM Tris-HCl, pH 7.5, 100 mM KCl, 5 mM MgCl_2_, 1 mM DTT, 5% glycerol), before initiating the cleavage reaction by addition of plasmid DNA. Final reaction concentrations were 1.5 μM protein:RNA complex and 500 ng DNA target. Reactions proceeded at room temperature or 37 °C, and aliquots were removed at selected time points and quenched with 50 mM EDTA and by the addition of 14 μg Proteinase K for 20 min. Cleavage products were resolved by gel electrophoresis on 1% agarose gel stained with GelRed (Biotium) and visualized using a Sapphire Biomolecular imager (Azure Biosystems). To quantify the cleavage activities, each gel image was analyzed using the ImageJ software (*39*) and plotted in GraphPad Prism. The percentage of DNA cleaved was determined by dividing the amount of cleaved DNA by the sum of uncleaved and cleaved DNA. The sequences of DNA and RNA oligonucleotides used in the study are provided in **source data 2**.

### Fluorescence Polarization Assay

The fluorescence polarization assay was performed in a 96-well plate format using a total reaction volume of 50 μL. A 20-nM, Fluorescein-labelled 30 base-pair dsDNA containing on target and mismatches at PAM distal-end positions were titrated against increasing concentrations of the SpCas9:gRNA (1:1.2) complex in a 20 mM tris(hydroxymethyl)-aminomethane (Tris)-HCl buffer, 150 mM KCl, 5 mM MgCl_2_, 5 mM EDTA, 1 mM DTT, 0.05% Tween 20, pH 7.5. Fluorescence anisotropy was measured at 25 °C with excitation at 485 nm and emission at 525 nm. The fluorescence polarization signal was measured using a microplate reader (ID5, Molecular Devices). Error bars represent ± standard deviation (s.d.) across technical replicates (n = 3). The sequences of DNA and RNA oligonucleotides used in the study are provided in **source data 2**.

## Supplementary files

Source data 1

Source data 2

Source data 3

## Acknowledgments

This research was supported by grants from NIAID (R01AI129719) and NIGMS (R35GM125034) to N.G.S. H.A. acknowledges funding from NIH (GM136859) and the Claudia Adams Barr Program for Innovative Cancer Research. The authors would like to acknowledge Dr. Kushol Gupta for assistance with SEC-MALS collection and analysis, and Dr. Ross Wilson for sharing the plasmid for sgRNA in vitro transcription.

## Author contributions

V.S.P. and N.G.S. designed the study and wrote the manuscript. V.S.P. generated constructs, prepared and purified isotopically labeled proteins and performed in vitro experiments and functional assays. V.S.P., A.D. and H.A. acquired and analyzed NMR data.

---

**Figure 1– figure supplement 1.**
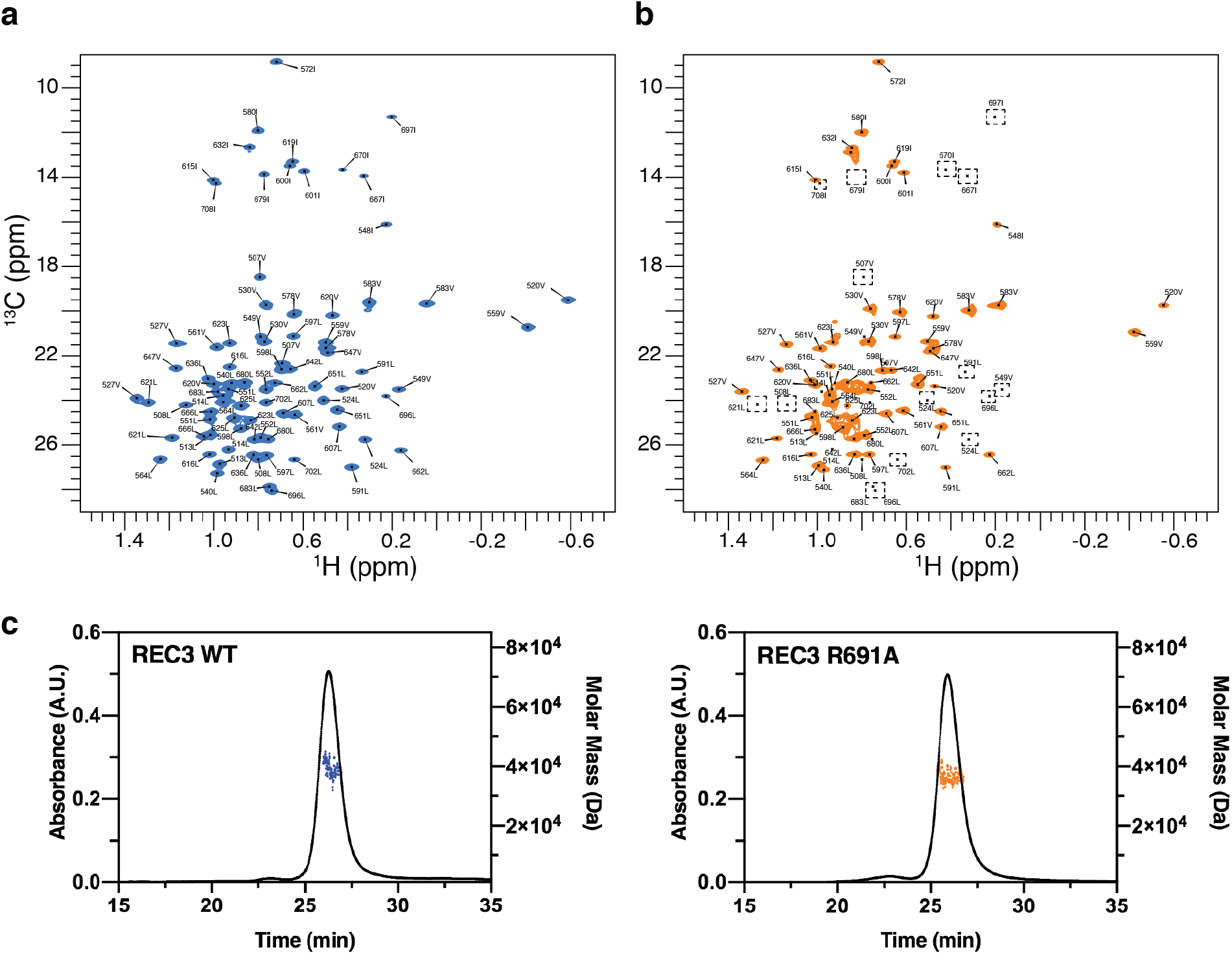
NMR spectra and characterization of the WT and R691 REC3 domain. ^1^H-^13^C methyl HMQC spectra of [U-^2^H,^15^N, Ileδ_1_-^13^CH_3_; Leu, Val-^13^CH_3_/^13^CH_3_] acquired at 800 MHz, 25 °C for **(a)** WT REC3 and **(b)** R691A REC3. Dotted black boxes highlight methyl resonances that become exchange broadened in the R691A REC3 mutant. **(c)** SEC-MALS of WT and R691A REC3 show that the proteins are monodispersed in solution.

**Figure 2– figure supplement 1.**
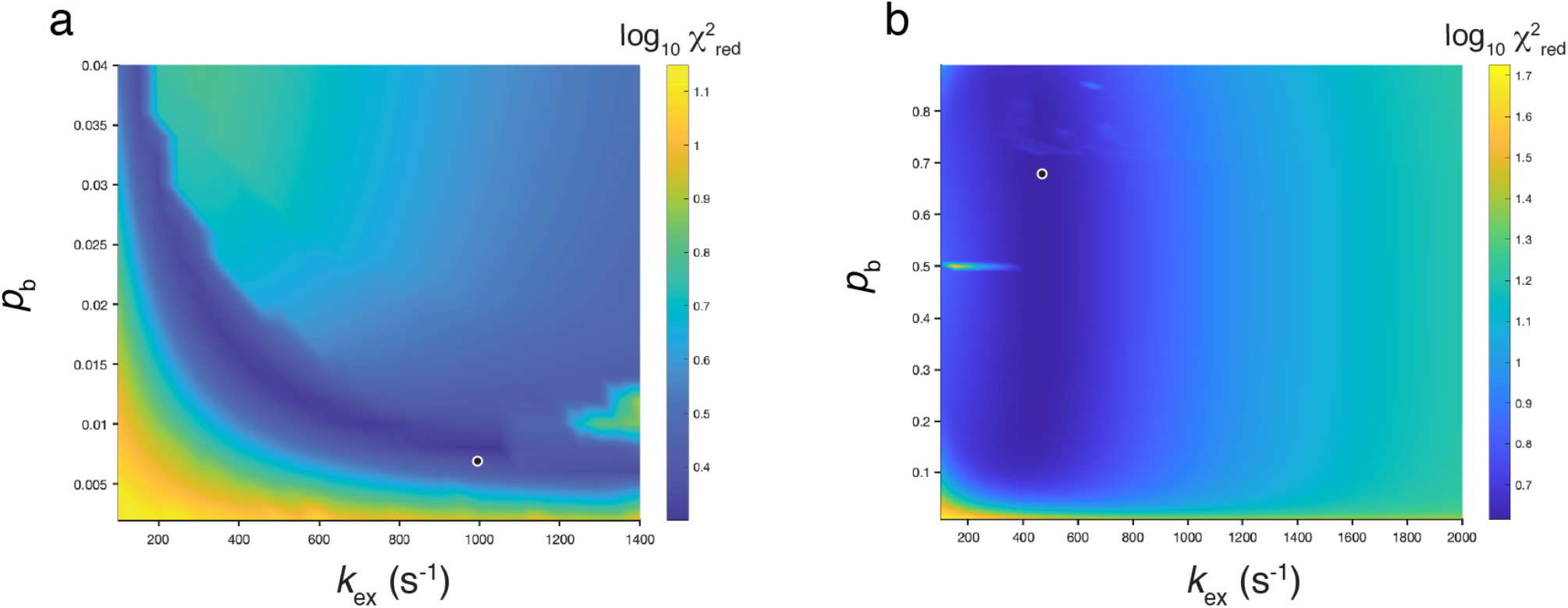
Reduced χ^2^ surfaces as a function of *pb* and *k*_*ex*_ obtained from a global analysis of the SQ ^13^C-CPMG data. χ^2^ surface plots (shown as log_10_) generated from fits of CPMG relaxation dispersion data recorded at 600 and 800 MHz on ILV-methyl labeled REC3 samples, WT (**a**) and R691A mutant (**b**). The |Δω| values were not fixed during the search for the minimum χ^2^ value. The white circle indicates the position of (*p*_*b*_, *k*_ex_) that corresponds to the global minimum of χ^2^. Values of (*p*_b_, *k*_ex_)=(0.8 ±0.05%, 988 ±88s^−1^), (68 ±14%, 468 ±19s^−1^) were obtained in **a** and **b**, corresponding to χ^2^ values of 0.30 and 0.62, respectively.

**Figure 2– figure supplement 2.**
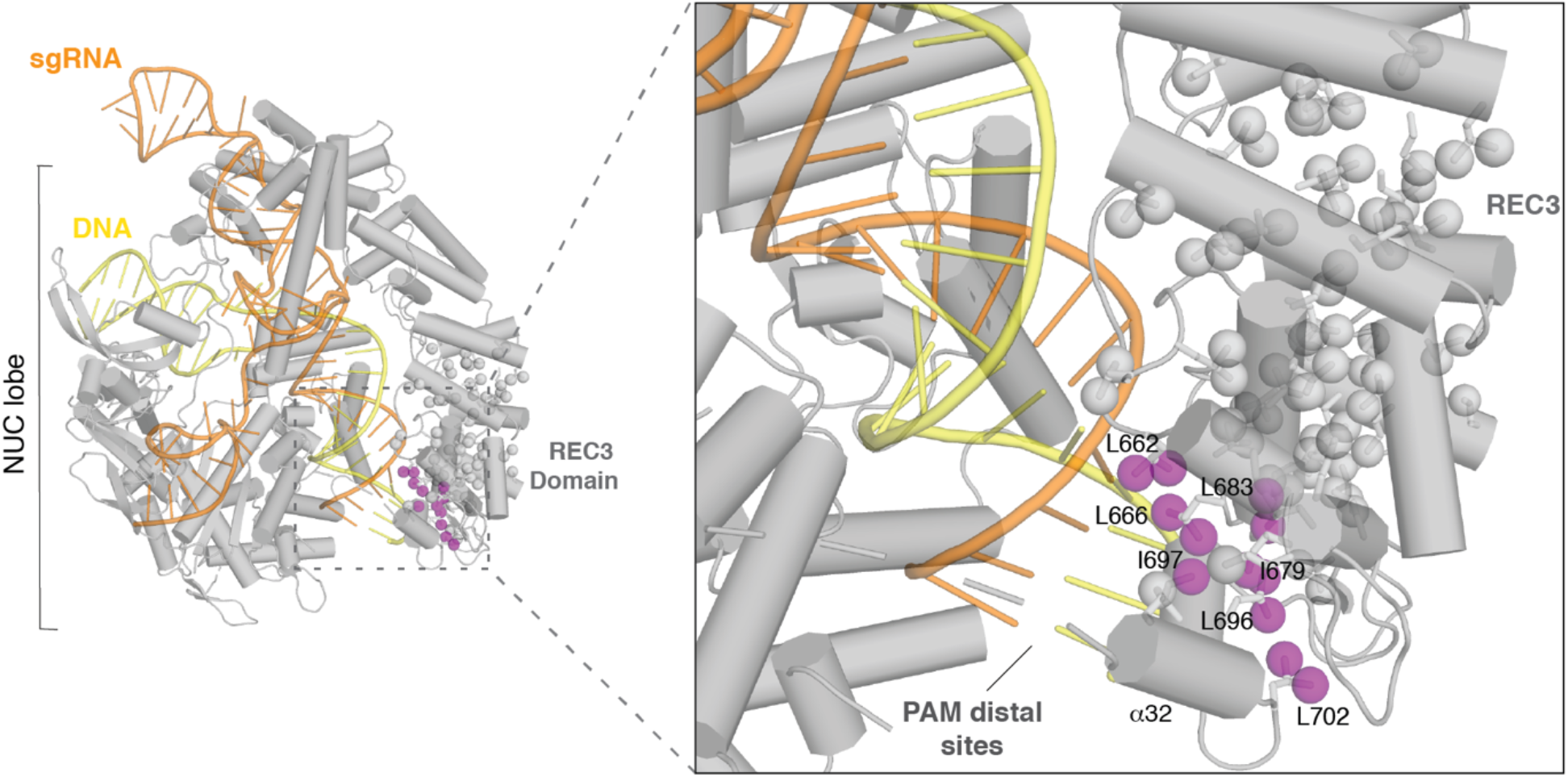
Local conformational plasticity in the α30-α32 helical motif of the REC3 domain. Crystal structure of sgRNA/DNA-bound Cas9 (Protein Data Bank (PDB) accession number 4UN3) with a detailed view of the DNA-RNA heteroduplex interface on the α30-α32 helices region of the REC3 domain. Methyl probes undergoing chemical exchange by single quantum ^13^C-CPMG are shown as purple spheres on the REC3 domain. The base pair mismatches associated with off-target effects, at PAM distal sites, are indicated.

**Figure 3– figure supplement 1.**
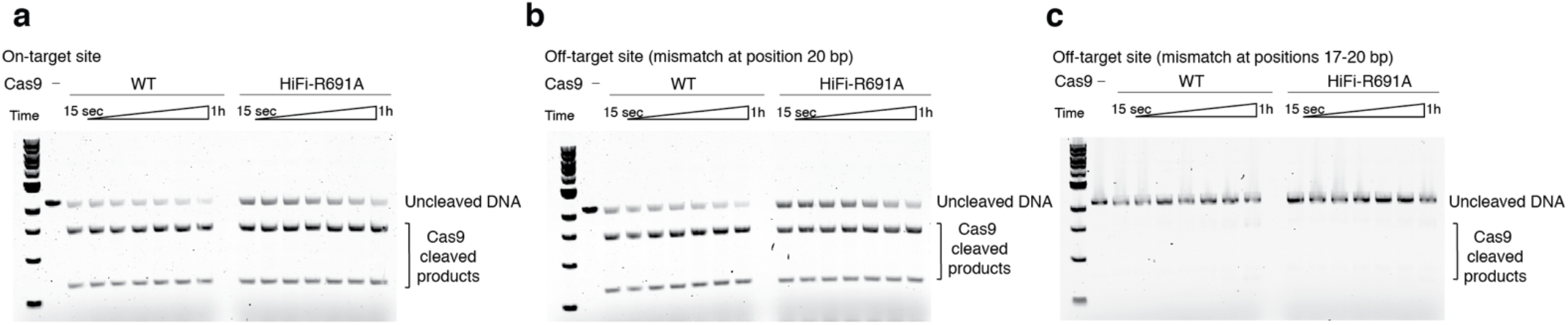
Endonuclease activities of Cas9 proteins containing mutations in the PAM-distal end. Endonuclease activity assays of wild type (WT) and R691A HiFi Cas9 proteins using a linearized plasmid template comprising a 20-base pair *EMX1* site containing a target sequence fully complementary to the sgRNA **(a)**, a target sequence mismatched to the sgRNA at position 20 **(b)**, or target sequence mismatched to the sgRNA at positions 17-20 **(c)**.

**Figure 4– figure supplement 1.**
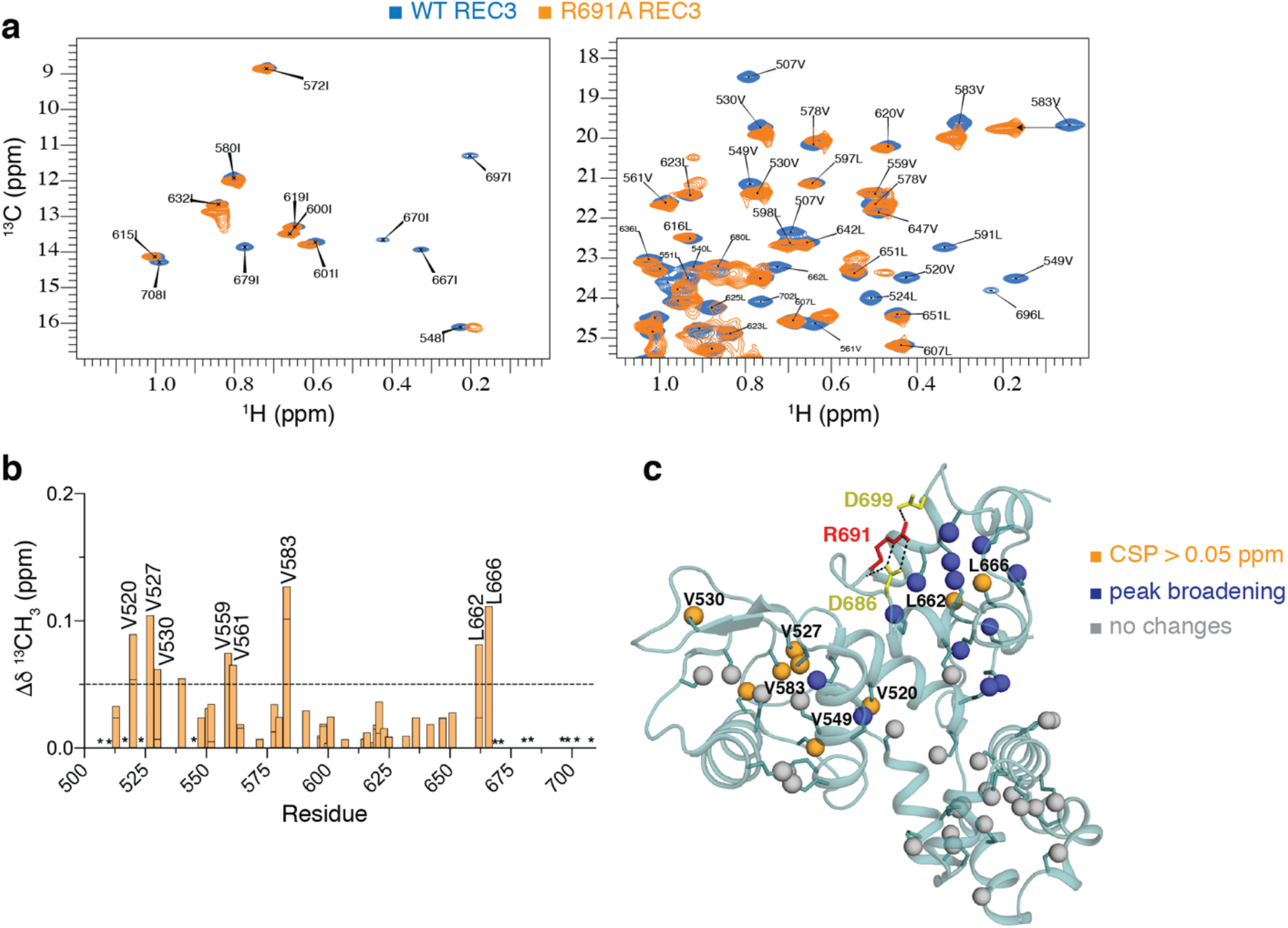
NMR chemical shift perturbations caused by the high-fidelity R691A mutation of REC3 domain. **(a)** Superposition of the ^1^H-^13^C-HMQC spectra of ILV-methyl labeled WT (blue) and R691A (orange) REC3 shown for the Ile and Leu/Val spectral regions on left and right panels, respectively. Assignments are indicated for residues with the largest chemical shift changes. Data acquired at 800 MHz, 25 °C. **(b)** Histograms of chemical shift perturbations in ILV-methyl labeled REC3. Residues with CSPs values 1 s.d. above the average are indicated (black dotted line). CSPs were calculated as described in Materials and Methods. Residues with line broadening beyond the detection are indicated with an asterisk. **(c)** Mapping of the methyl groups with marked chemical shift differences (orange spheres) and peak broadening (blue spheres) onto the REC3 structure. The results of the effect of this mutation on the dynamics of REC3 are shown in **Figure 4**. The change in chemical shift (in p.p.m.) between the WT and R691A REC3 was determined using the equation Δδ^CH3^ = [1/2 (Δδ^2^H + Δδ^2^C/4)]^1/2^.

**Figure 4– figure supplement 2.**
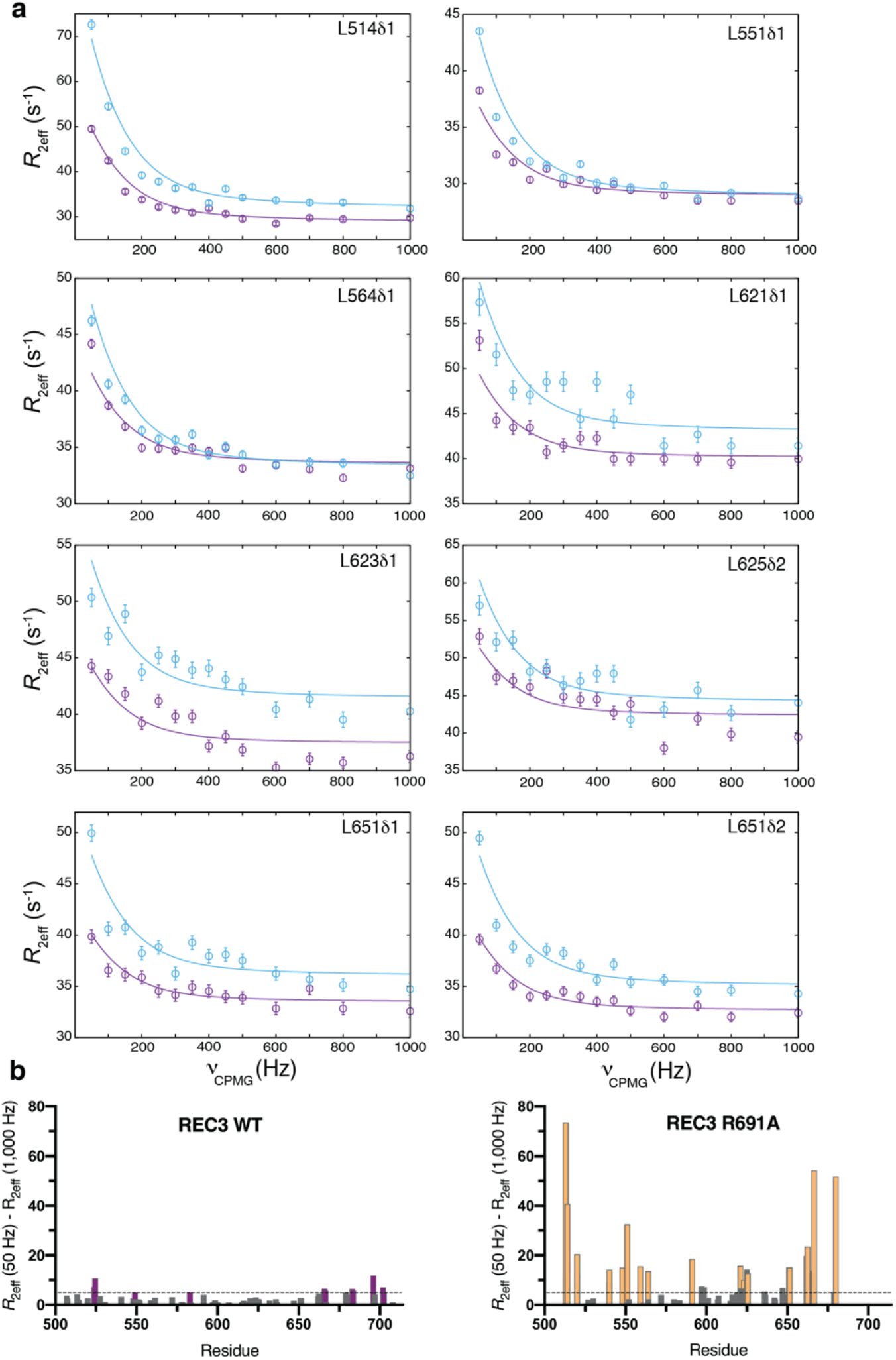
Relaxation dispersion experiments indicate that R691A mutant increases conformational exchange in HiFi REC3 domain. **(a)** Experimental relaxation dispersion profiles (circles) for residues exhibiting μs-ms timescale dynamics as measured by ^13^C SQ CPMG relaxation dispersion experiments for R691A REC3, recorded at 15°C (800 MHz – blue; 600 MHz – purple). Solid lines represent the best fit to a global two-site exchange model. Error bars indicate uncertainties in *R*_2eff_ rates. **(b)** Plots of the *R*_ex_ contributions of the methyl groups for WT and R691A mutant. *R*_ex_ contributions were calculated from the differences between R_2eff_ (50 Hz) and R_2eff_ (1000 Hz). The methyl groups with significant *R*_ex_ contributions (> 5 Hz) are colored purple for WT and orange for R691A REC3 domain.

**Figure 4– figure supplement 3.**
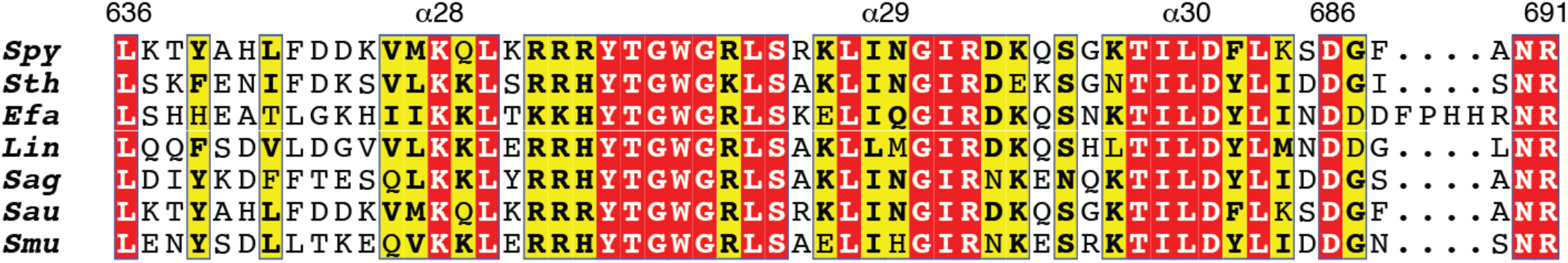
Multiple sequence alignment of Type II-A Cas9 orthologs focusing on the α28-α31 helices region. The primary sequences of Type II-A Cas9 orthologs from *Streptococcus pyogenes* (Spy), *Streptococcus thermophilus* (Sth), *Listeria innocua* (Lin), *Streptococcus agalactiae* (Sag), *Streptococcus mutans* (Smu), *Enterococcus faecium* (Efa) and *Staphylococcus aureus* (Sau) were aligned using Clustal Omega. The alignment was illustrated by ESPscript with default settings.

